# Genome-enabled inference of functional genetic variants in the face, brain and behavior

**DOI:** 10.1101/2020.10.12.336453

**Authors:** Chinar Patil, Jonathan B. Sylvester, Kawther Abdilleh, Michael W. Norsworthy, Karen Pottin, Milan Malinsky, Ryan F. Bloomquist, Zachary V. Johnson, Patrick T. McGrath, Jeffrey T. Streelman

## Abstract

Lake Malawi cichlid fishes exhibit extensive divergence in form and function among closely related species separated by a relatively small number of genetic changes. During the past million years, hundreds of species have diversified along an ecological axis in rock vs. sand habitats. We compared the genomes of rock- and sand-dwelling species and asked which genetic variants in which genes differed among the groups. We found that 96% of differentiated variants reside in non-coding sequence but these non-coding diverged variants are evolutionarily conserved. The majority of divergent coding variants are missense and/or loss of function. Regions near differentiated variants are enriched for craniofacial, neural and behavioral functional categories. To follow up experimentally, we used rock- vs. sand-species and their hybrids to (i) clarify the push-pull roles of BMP signaling and *irx1b* in the specification of forebrain territories during gastrulation and (ii) reveal striking context-dependent brain gene expression during adult social behavior. Our results suggest compelling ties between early brain development and adult behavior and highlight the promise of evolutionary reverse genetics – the identification of functional variants from genome sequencing in natural populations.

## Introduction

Our understanding of how the genome encodes natural variation in form and function is still limited. This is the case for almost any trait, from height to behavior to complex disease (Boyle, Li et al. 2017). The reasons for this are manifold, but they include an underappreciated role of non-coding genetic variants linked to differences in traits. This is apparent in our assumptions and in syntheses of data. For instance, only 25 years ago, experts thought that the human genome might contain 70,000 to over 100,000 genes to account for our complexity (Fields, Adams et al. 1994). More recently, it has been estimated that upwards of 93% of human disease related variants – traits for which we have the most data from genome wide association studies (GWAS) – reside in noncoding DNA sequence (Maurano, Humbert et al. 2012). Many of these noncoding variants are regulatory, that is, they affect the expression of genes (Degner, Pai et al. 2012). Therefore, despite a refined understanding of how single genes work in controlled cellular environments, it remains unclear how the genome is activated to produce natural phenotypes, and this may be particularly vexing for context-dependent processes like development or behavior.

Over the past two decades, systems have been developed to identify the genetic basis of traits from nature(Streelman, Peichel et al. 2007). Amongst vertebrate animals, these traits include body armor (Colosimo, Peichel et al. 2004), color (Kratochwil, Liang et al. 2018), head and jaw shape (Albertson, Streelman et al. 2005, Shapiro, Kronenberg et al. 2013, Lamichhaney, Berglund et al. 2015), parental care (Okhovat, Berrio et al. 2015, Bendesky, Kwon et al. 2017), song (Pfenning, Hara et al. 2014) and coordinated movement (Greenwood, Wark et al. 2013). The take home message from this work has been that a small number of genes from recognizable pathways explain a considerable proportion of phenotypic variance. Yet, these studies may be biased in interpretation and limited in inference space. The focus is typically on one or two species and one trait at a time, often using hybrid pedigrees founded by a small number of individuals, and candidate gene or QTL approaches. Here, we explored a different strategy in a system of many species with many divergent traits. We sought to determine the genetic differences between closely related groups of species and then to focus experiments on leads from genome divergence. In essence, we’ve asked the genome which traits to follow.

The Malawi cichlid system is an apposite one for our research aims. The assemblage comprises hundreds of closely related species that have diversified in the last 500,000 to one million years (Kocher 2004), such that the genomes of individuals across species boundaries remain highly similar (Loh, Katz et al. 2008). An appreciable fraction of genetic polymorphism identified in Malawi species is shared with cichlid lineages from throughout East Africa -- suggesting that ancient genetic variation fuels diversification of the Malawi flock (Loh, Bezault et al. 2013). Set against this background of genome similarity, Malawi cichlids exhibit staggering diversity in phenotypes including pigmentation (Streelman, Albertson et al. 2003), sex determination (Roberts, Ser et al. 2009, Parnell and Streelman 2013), craniofacial and brain patterning (Albertson, Streelman et al. 2005, Sylvester, Rich et al. 2010, Sylvester, Rich et al. 2013) and social behavior (York, Patil et al. 2018, Baran and Streelman 2020, Johnson, Moore et al. 2020). Previous work has focused on the genomic and early developmental underpinnings of this diversity, in rock- *vs*. sand-dwelling species (Loh, Katz et al. 2008, Fraser, Hulsey et al. 2009, Sylvester, Rich et al. 2010, Sylvester, Rich et al. 2013).

Rock- vs. sand-species form ecologically distinct groups similar to other ecotypes in well-known adaptive radiations (marine *vs*. freshwater sticklebacks; tree *vs*. ground finches and anoles) (Streelman and Danley 2003). The main difference in this case is that each of the rock- and sand-groups contains more than 200 species. Recent divergence, rapid speciation and meta-population dynamics synergistically lead to the broad sharing of polymorphism across the rock-sand speciation continuum (Loh, Bezault et al. 2013, Malinsky, Svardal et al. 2018). Malawi rock-dwellers tend to be strongly territorial and aggressive; they breed and feed at high density in complex rock-reef habitats. Most eat algae from the substratum with strongly reinforced jaws packed with teeth. Adult rock-dweller brains exhibit enlarged anterior components, telencephala and olfactory bulbs. Sand-dwellers are less site-specific and less aggressive. They often breed on communal leks where males build sand ‘bowers’ to attract females (McKaye, Louda et al. 1990). Many capture small prey using acute vision and fast-moving gracile jaws; their brains and sensory apparatus are elaborated for more posterior structures optic tecta, thalamus and eyes (SI Figure 1). We aimed to understand evolutionary divergence between rock- and sand-dwelling lineages by identifying the number, type and spectrum of genetic variants that separate these groups.

To target this primary axis of evolutionary divergence in the Lake Malawi species assemblage (Streelman and Danley 2003), we compared whole genomes of one male individual each from 8 rock-dwelling and from 14 sand-dwelling species (SI Table 1), to an average of 25X coverage per individual. Species were chosen to represent the diversity present within each of the rock- and sand-groups (Figure 1A), in terms of body size, color pattern, ecology and phylogenetically defined lineages within the sand-species group (Malinsky, Svardal et al. 2018).

**Table 1.**

| Species | Habitat | Sample Source | Accession number | UMD2a_map_percent | UMD2a_meanco v | UMD1_map_percent | UMD1_meanco v |
| --- | --- | --- | --- | --- | --- | --- | --- |
| *Metriaclima zebra* | Rock | Lab bred | SRR63 22515 | 97.50 | 42.63 | 96.81 | 46.68 |
| *Labeotropheus trewavasae* | Rock | Streelman Finclip #2238 | SRR63 14224 | 98.23 | 14.89 | 97.53 | 16.42 |
| *Labeotropheus fuelleborni* | Rock | Lab bred | SRR63 14225 | 97.78 | 27.02 | 97.09 | 29.79 |
| *Cynotilapia afra* | Rock | Lab bred | SRR63 14226 | 98.89 | 23.88 | 98.24 | 26.33 |
| *Cyathochromis obliquidens* | Rock | Streelman Finclip #2368 | SRR63 14227 | 98.52 | 13.95 | 97.82 | 15.38 |
| *Metriaclima patricki* | Rock | Lab bred | SRR63 14228 | 97.78 | 37.41 | 97.20 | 41.21 |
| *Petrotilapia 'chitimba'* | Rock | Lab bred | SRR63 14229 | 95.04 | 27.82 | 94.20 | 30.60 |
| *Genyochromis mento* | Rock | Streelman Finclip #2170 | SRR63 14230 | 91.77 | 8.83 | 91.10 | 9.74 |
| *Mchenga conophorus* | Sand | Lab bred | SRR54 38125 | 98.39 | 35.44 | 97.70 | 39.05 |
| *Ctenopharynx nitidus* | Sand | Martin Genner [#17] | SRR54 38124 | 97.21 | 30.22 | 96.39 | 33.26 |
| *Copadichromis likomae* | Sand | Martin Genner [#52] | SRR54 38110 | 96.72 | 27.06 | 96.08 | 29.83 |
| *Copadichromis* sp.<br>"mloto goldcrest"<br>*sensu Konings* | Sand | Lab bred | SRR54<br>38123 | 99.20 | 21.68 | 98.57 | 23.93 |
| *Tramitichromis*<br>"chembe" | Sand | Ryan<br>York | SRR54<br>38109 | 87.84 | 16.50 | 87.28 | 18.20 |
| *Taeniolethrinops*<br>*furcicauda* | Sand | Martin<br>Genner<br>[#25] | SRR54<br>38122 | 87.54 | 15.48 | 86.96 | 17.07 |
| *Tramitichromis*<br>*intermedius* | Sand | Lab bred | SRR54<br>38118 | 97.81 | 35.60 | 97.04 | 39.29 |
| *Copadichromis*<br>*virginalis* "yellow<br>blaze nkanda" | Sand | Lab bred | SRR54<br>38119 | 97.96 | 34.54 | 97.28 | 38.07 |
| *Fossochromis*<br>*rostratus* | Sand | Streelma<br>n finclip<br>stock<br>[2549] | SRR54<br>38116 | 97.32 | 28.95 | 96.77 | 31.97 |
| *Aulonocara*<br>*baenschi* | Sand | Lab Bred | SRR54<br>38115 | 89.31 | 28.04 | 88.71 | 30.91 |
| *Dimidiochromis*<br>*compressiceps* | Sand | Lab Bred | SRR54<br>38114 | 95.78 | 24.62 | 95.18 | 27.16 |
| *Mylochromis*<br>*sphaerodon* | Sand | Lab Bred | SRR54<br>38113 | 97.37 | 24.63 | 96.60 | 27.12 |
| *Nimbochromis*<br>*polystigma* | Sand | Ryan<br>York | SRR54<br>38113 | 81.57 | 16.77 | 81.06 | 18.49 |
| *Dimidiochromis*<br>*kiwinge* | Sand | Streelma<br>n finclip<br>stock<br>[2384] | SRR54<br>38112 | 96.46 | 16.33 | 95.93 | 18.03 |

**Figure 1:**
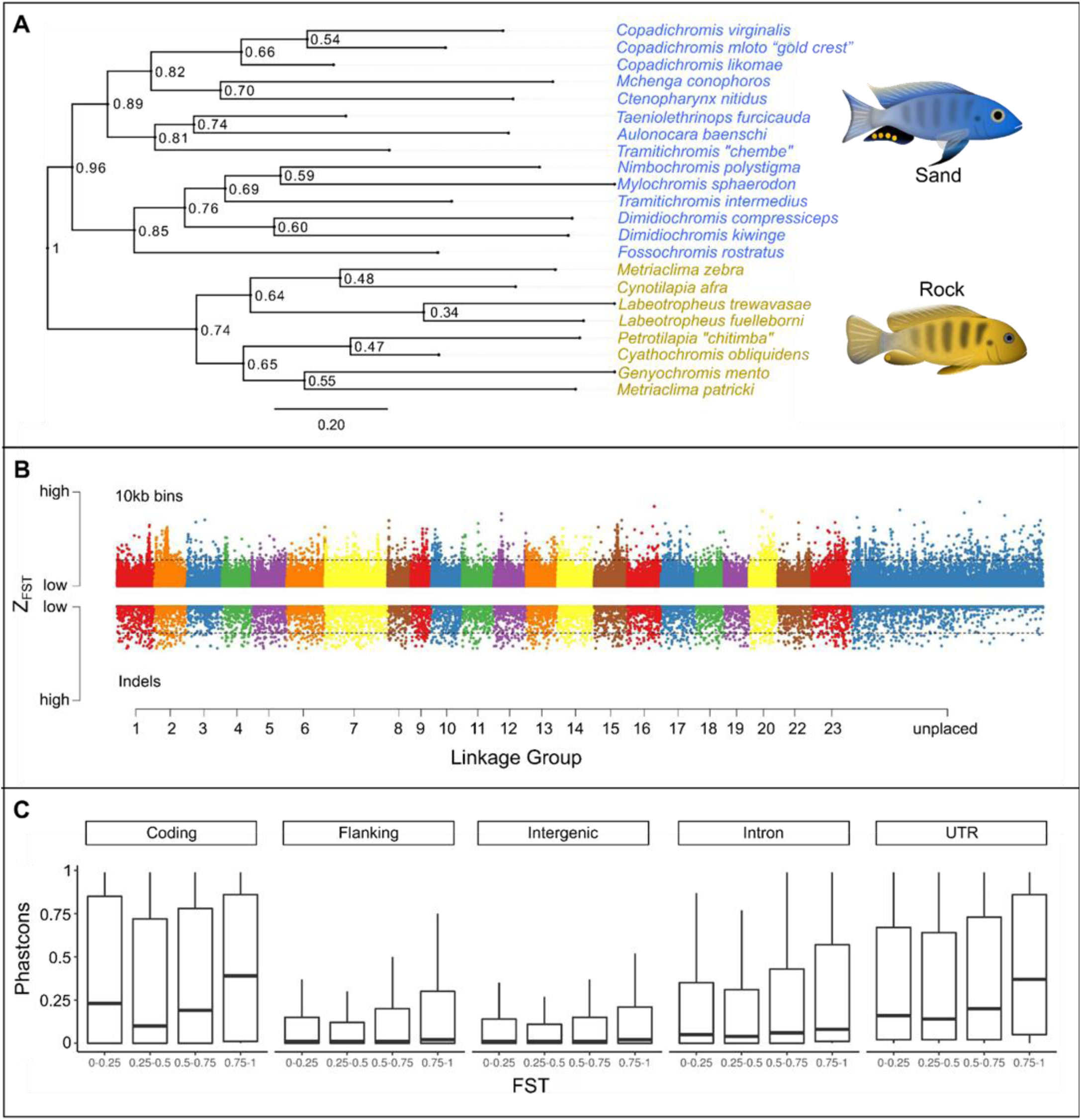
The genomic substrate for rock vs. sand evolution. (A) A maximum likelihood phylogeny of eight rock- and fourteen sand-dwelling species, based on variable sites (informative SNP and InDels) identified throughout the genome. “Sand” species contain representatives of the lineages “shallow benthic, deep benthic and utaka” from (Malinsky, Svardal et al. 2018), while the “rock” species correspond to the “mbuna” lineage. (B) A plot of Z-F_ST_ (F_ST_ normalized using Fisher’s Z-transformation) across the genome, plotting genomic divergence between rock- vs. sand-dwelling groups. Single nucleotide polymorphisms (SNPs) summed over 10kb bins and insertion-deletion mutations (InDels) are shown on the same scale. Numbers along the x-axis refer to linkage groups (i.e., chromosomes) and threshold lines indicate 2.5% FDR. (C) Evolutionary conservation (PhastCons) scores were calculated for each nucleotide across the genome, subdivided by genome annotation and plotted by bins of increasing F_ST_. The PhastCons score for each genome category is significantly higher for increasing bins of F_ST_ (Wilcoxon rank sum p value < 2e-16).

## Results

### The genomic signature of rock-sand divergence

We compared the genomes of 8 rock dwellers and 14 sand dwellers to uncover the genomic signature of rock-versus sand-evolutionary diversification. We aligned sequence data to a reference genome of nearly 1 gigabase (Conte, Joshi et al. 2019) and identified approximately 22 million Single Nucleotide Polymorphisms (SNPs) and 200,000 Insertion-Deletions (InDels). We calculated F_ST_ per variant, and averaged across 10kb windows, to quantify divergence between rock and sand species. We found that 0.06% of SNPs and 0.44% of InDels are alternately fixed between rock- and sand-groups. When these divergent variants and genome regions (2.5% FDR) are mapped to linkage groups (chromosomes), it is apparent that the signature of rock- vs. sand-divergence is distributed relatively evenly across the chromosomes (Figure 1B). Among fixed variants, 3.5% were found in coding regions and 96.5% were predicted to be non-coding; ∼17% in intergenic regions, 38% in introns, 38% in flanking regions (within 25kb up- or downstream of a gene), and 3% in annotated UTRs. Rock vs. sand fixed coding variants were more likely to be missense/loss-of-function (72.6%) than silent (27.3%).

We next generated whole-genome alignments of five published cichlid reference genomes from across East Africa (Brawand, Wagner et al. 2014) and estimated an evolutionary conservation score for each nucleotide position. Akin to phylogenetic footprinting, this approach allows inference of function for regions that are slower to change than others due to the long-term effect of purifying selection. For both coding and non-coding portions of the genome, we found that rock-sand divergence correlates positively with evolutionary conservation scores (Figure 1C), suggesting that differentiated rock-sand variants, including many non-coding variants, are enriched for function.

A total of 4,484 genes lie within 25 kb of either an alternately fixed variant or a highly divergent 10kb window (2.5%FDR). Pathway enrichment analysis (Ben-Ari Fuchs, Lieder et al. 2016) of human homologs/analogs for these genes reveals categories spanning early embryonic development, craniofacial morphogenesis, brain development, synaptic transmission and neuronal function (SI Table 2). In particular, rock-sand divergent genes are enriched for GO Biological Process terms ‘telencephalon development’ (p < 1.7e-18), ‘adult behavior’ (p < 2e-14), ‘synaptic plasticity’ (p < 1.4e-12), ‘odontogenesis’ (p < 3.7e-11), ‘response to BMP’ (p < 3.2e-09), ‘gastrulation’ (p < 5.6e-06), ‘face morphogenesis’ (p < 8.9e-08), ‘neural crest cell differentiation’ (p < 4.3e-13), and ‘eye development’ (p < 1.3e-15). Over-represented gene families included nuclear hormone receptors (p < 3.0e-08), HOXL subclass homeoboxes (p < 1.4e-07), TALE class homeoboxes (p < 4.2e-04) and Forkhead boxes (p < 9.33-04; for novel expression domains in cichlid *foxp2* see SI Figure 2). We observed enrichment for the mouse phenotypes ‘abnormal cognition’ (p < 3.3e-15), ‘abnormal learning and memory’ (p < 2.6e-15), ‘abnormal craniofacial morphology’ (p < 4.9e-11) and ‘abnormal social/conspecific interaction’ (p < 7.1e-14). We used the list of differentiated genes to query an Allen Brain Atlas dataset that reports gene expression in hundreds of brain regions (Hawrylycz, Lein et al. 2012). Rock-sand-divergent genes were enriched for the basomedial nucleus of the amygdala, a sub-region of the telencephalon (p_adj_ = 0.001) that regulates fear, anxiety, and physiological responses to territorial intruders in rodents (Adhikari, Lerner et al. 2015, Mesquita, Abreu et al. 2016), and has been linked to Social Anxiety Disorder in humans (Carvalho, Nóbrega et al. 2020). Finally, we queried databases of genes involved in human disease. Genes near divergent variants are significantly enriched for factors implicated in neurological disease like Autism Spectrum Disorder (SFARI (Abrahams, Arking et al. 2013), Fisher’s exact test p value < 2e-16) and disorders related to the neural crest (Piñero, Bravo et al. 2017), (Fisher’s exact test p value < 2e-16).

**Figure 2:**
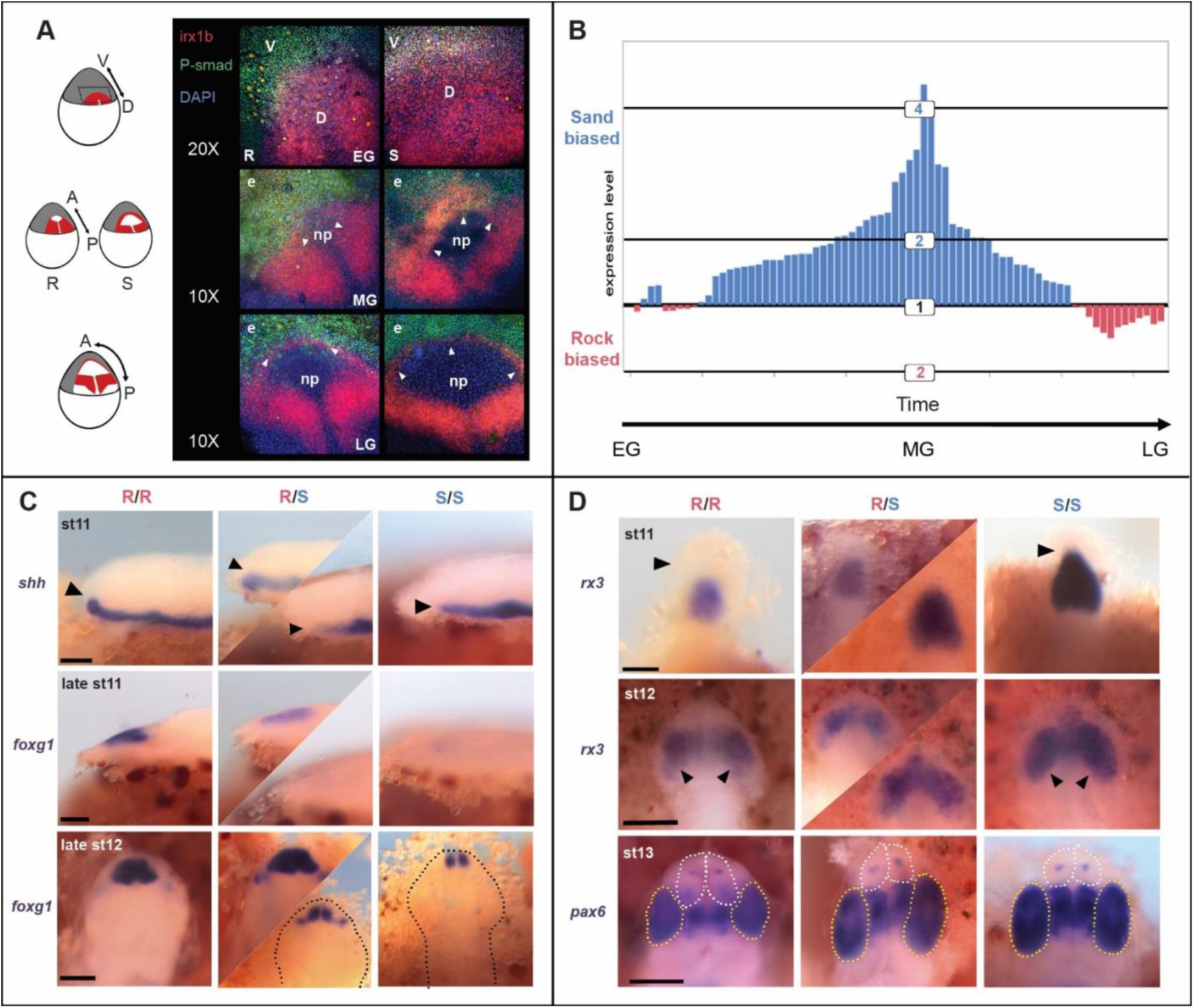
A gastrula-stage map of forebrain diversification. (A) Double in situ hybridization – immunohistochemistry to visualize *irx1b* expression (red) and BMP [PSMAD] activity (green) across the stages of cichlid gastrulation; DAPI in blue. Three rows represent early, mid and late gastrulation (EG, MG, LG) in embryos of rock- (R) and sand-dwelling (S) species. Sand-dwellers show expanded *irx1* expression in the dorsal portion of the embryo at EG, expanded *irx1b* expression in the anterior domain at MG (arrowheads) and clear PSMAD activity from the developing neural plate (np) earlier in LG (arrowheads). e=epidermis. Schematics at left show *irx1b* expression domains in red, on cartoons of cichlid embryos. (B) Relative expression of rock- (red) and sand- (blue) *irx1b* alleles, sampled from 74 heterozygous rock X sand F_2_ embryos, across the stages of gastrulation. F_2_ embryos were sampled at stage 9 (gastrulation). Because Malawi cichlid species are maternal mouthbrooders and eggs are fertilized in batches per brood, each brood’s embryos vary in timing of fertilization by up to 4h. Embryos within broods can therefore be sub-staged in gastrulation (see methods). Each bar on the plot represents the relative allelic expression of sand- and rock-*irx1b* in a heterozygous F_2_ individual. Quantification of allele-specific expression (ASE) shows that levels are sand-biased, and that this effect is strongest in MG. (C) in situ hybridization of *shh* and *foxg1*, during neurula stages, showing development of the telencephalon in rock-X sand-F_2_ embryos, indexed for *irx1b* genotype. F_2_ individuals homozygous for rock-*irx1b* alleles (R/R) show a more dorsal progression of *shh* expression (black arrowheads), an earlier and a larger expression domain of the telencephalon marker *foxg1*. The top two rows are lateral views; bottom row is a dorsal view. Dotted lines demarcate the outline of the embryo in dorsal view. Heterozygous individuals exhibit greater variation in expression domains (middle columns), indicating that genetic factors other than variants in *irx1b* contribute to this phenotype. (D) in situ hybridization for *rx3* and *pax6*, during neurula stages, chart the development of the eye field in rock-X sand-F_2_ embryos indexed for *irx1b* genotypes. F_2_ individuals homozygous for sand-*irx1b* alleles (S/S) show larger domains of *rx3* (black arrowheads) and larger eyes (*pax6*, also marked by yellow dotted line), but smaller telencephala (white dotted line). All panels are dorsal views. Heterozygous individuals exhibit greater variation in expression domains (middle columns), indicating that genetic factors other than variants in *irx1b* contribute to this phenotype.

Given the prevalence of evolutionarily conserved, non-coding, divergent rock-sand variants and genome-wide enrichment for craniofacial and neural crest biology, we examined overlap with published datasets of mammalian neural crest and craniofacial enhancers (Rada-Iglesias, Bajpai et al. 2012, Attanasio, Nord et al. 2013). These data allow us to identify craniofacial and cranial neural crest cell (CNCC) enhancers conserved between mammals and cichlids and fixed variants between rock and sand species within these conserved regulatory elements. A total of 275 craniofacial enhancer elements and 234 human CNCC enhancers are evolutionarily conserved between mammals and cichlids. We found divergent rock-sand mutations within the enhancer elements of key genes integral to CNCC specification and migration (SI Table 2). Notably, from both datasets, fixed rock-sand variants were found within the enhancer region of the gene *nr2f2*, a nuclear receptor and master neural crest regulator (Simoes-Costa and Bronner 2015). Rock-sand divergent variants were similarly located within craniofacial enhancers of three genes (*yap1, fat4, rere*) that function in the Hippo signaling pathway, as well as within enhancers of *irx3* and *axin2*. These data linking rock-sand fixed SNP/InDels to evolutionarily conserved, experimentally verified enhancers further underscore the importance of non-coding variation in the craniofacial evolution of rock- and sand-lineages (Roberts, Hu et al. 2011).

Genome-wide divergence between rock vs. sand Malawi cichlids involves a relatively small percentage of genetic variants. Divergent variants are (a) predominantly non-coding, (b) in long-term evolutionarily conserved loci (c) enriched for genes and pathways involved in embryonic development, brain development, brain function and behavior, and craniofacial morphogenesis. Given these strong patterns of enrichment, we used the experimental power of the Malawi cichlid system to interrogate features of early development and adult behavior that differ between rock- and sand-groups.

### A gastrula-stage map of forebrain diversification

Rock- vs. sand-dwelling Malawi cichlids exhibit divergence in or near genes enriched for BMP signalling, gastrulation, eye and telencephalon development, as well as the TALE (Irx) gene family. To explore the developmental consequences of this differentiation, we investigated early forebrain specification in rock- and sand-embryos, building upon our previous studies and interest in *irx1b* and early brain development (Sylvester, Rich et al. 2010, Sylvester, Rich et al. 2013). During development, the complexity of the vertebrate brain is first laid out in the neural plate, a single-cell thick sheet of cells that forms between non-neural ectoderm and the germ ring at gastrulation. Irx genes act as transcriptional repressors of BMP signal in gastrulation, and function to specify the neural plate (Cavodeassi, Modolell et al. 2001). BMPs, in turn, are protective of the anterior-most region of the neural plate, which will ultimately give rise to the telencephalon, and suppress the eye field (Bielen and Houart 2012). Given alternatively fixed variants in the *irx1b* gene, expected interactions between Irx and BMP signaling in the early embryo and known telencephalon vs. eye size differences between rock- vs. sand-species (Sylvester, Rich et al. 2010, Sylvester, Rich et al. 2013), we examined and quantified the early activity of *irx1b* and BMP in rock- vs. sand-embryos.

We used a custom device to orient and image cichlid embryos *in toto* at gastrula and neurula stages (White, Sylvester et al. 2015). In early gastrula (EG), *irx1b* (red) and BMP signal (green, PSMAD) delineate complementary dorsal and ventral domains of the embryo (Figure 2A). By mid-gastrula (MG), *irx1b* shows two expression domains, one in the posterior portion of the developing neural plate (np) and the second co-expressed with PSMAD activity around its anterior border (white arrowheads). By late gastrula (LG), the domains of *irx1b* expression and PSMAD activity sharpen around the leading edge of the neural plate but remain overlapping around the periphery. Notably, *irx1b* expression is expanded in the anterior domain of sand-dwellers (S) compared to rock-dwellers at EG and MG, and then defines the boundary of the neural plate earlier in sand-dwellers (S) in LG (arrowheads). As a consequence, BMP signal should have a longer-lasting influence on the neural plate in rock-dwelling species. Based upon manipulative experiments in zebrafish (Bielen and Houart 2012), this is predicted to result in a relatively larger presumptive telencephalon and smaller eye field.

We developed a panel of rock-x sand-hybrid crosses to formally evaluate the role of *irx1b* in forebrain diversification. First, we used quantitative RT-PCR to measure allele-specific expression (ASE) in heterozygous rock-x sand-F_2_ hybrids, across the stages of gastrulation. We observed that the sand-*irx1b* allele was expressed at significantly higher levels (average of 2.5-fold; p = 4.5e-13; Student’s t-test) and that this difference was largely confined to MG (Figure 2B). Next, we used hybrid embryos to chart the development of the telencephalon and the eye field. Rock-x sand-F_2_ hybrids, indexed for *irx1b* genotype, were raised to neurula and somitogenesis stages and we examined the expression of *shh* (which induces *foxg1* and the ventral forebrain), *foxg1* (a marker of the telencephalon), and *rx3* (a marker of the eye field), by in situ hybridization. F_2_ individuals homozygous for rock-*irx1b* alleles exhibited a larger and more rostral domain of *shh* expression, an earlier and larger domain of *foxg1* and a smaller *rx3* domain (Figure 2C, D). These differences between rock- vs. sand-*irx1b* genotypes match expression divergence observed amongst rock- vs. sand-species (Sylvester, Rich et al. 2010, Sylvester, Rich et al. 2013). Finally, when we compared the relative size of the telencephalon among *irx1b* genotypes, individuals homozygous for rock-alleles exhibited larger telencephala (SI Figure 3). We conclude that genetic variants in and around the *irx1b* gene contribute to divergent specification of the Malawi cichlid forebrain, likely via spatial, temporal and quantitative variation in the expression of *irx1b* itself.

Our genome sequencing revealed a near-fixed InDel in the 3’ UTR of the Malawi cichlid *irx1b* gene (SI Figure 4). Rock-species possess an 85bp insertion, compared to cichlid species from outside of the Malawi lineage. Sand-dwellers largely lack the insertion and exhibit a 6bp deletion compared to outgroups. The insertion shows strong genetic similarity to a fragment from the Rex1 family of non-LTR retrotransposons (Volff, Korting et al. 2000). Given the likelihood that *Astatotilapia calliptera* populations surrounding Lake Malawi may have seeded the Malawi evolutionary radiation and contributed to rock- and sand-dwelling lineages (Loh, Bezault et al. 2013, Malinsky, Svardal et al. 2018), we explored the presence/absence of this InDel in *Astatotilapia* samples. We found that most *Astatotilapia* individuals and populations had the rock-*irx1b* allele (the insertion), but that an individual from Chizumulu Island was fixed for the sand-allele and two individuals sampled from Itupi were heterozygous. Because the 85bp insertion in rock-species is a partial Rex1 fragment, and sand-species carry a 6bp deletion compared to outgroups, we speculate that the current rock- and sand-divergent alleles were generated by at least two imperfect excision events of an element that invaded the genome of the Malawi + *Astatotilapia* ancestor. Rex1/Babar retrotransposons have been active in African cichlid genomes, and are known to influence gene expression when inserted in 5’ and 3’ UTRs (Brawand, Wagner et al. 2014). Future experiments will determine whether this Rex1 insertion causes the differences in *irx1b* gene expression and forebrain specification we observed.

### Genomics of divergent social challenge and opportunity

Rock- and sand-dwelling Malawi cichlids live in strikingly different social and physical environments. Rock-dwelling males tend to be more aggressive than sand-dwellers (Baran and Streelman 2020) and defend territories year-round as sites for feeding and breeding. By contrast, sand-dwellers are more exploratory than rock-dwellers (Johnson, Moore et al. 2020) and only breeding males tend to be territorial, often building sand bowers to attract females and mitigate male-male aggression. Given these observations and genome-wide enrichment for categories related to adult behavior and social interaction, we designed an experimental paradigm to investigate brain gene expression profiles associated with divergent rock- vs. sand-social behaviors.

We evaluated social challenge and opportunity amongst males using a large tank with a ‘rock’ habitat at one end and ‘sand’ at the other, separated by glass bottom (Figure 3A). When parental rock-species are placed in this tank paradigm, males court females on the rock side of the tank. Males of sand-species court females over sand and construct species-appropriate bowers. When single hybrid rock-x sand-F_1_ males are placed in this arena with hybrid F_1_ females, males invariably court females over the ‘rock’ habitat. However, when two rock-x sand-hybrid F_1_ males (brothers) were allowed to compete for gravid hybrid F_1_ females in this tank paradigm, we observed something different. One male, typically the larger, courted females over the rock habitat, and the other simultaneously constructed bowers to court females over the sand. We found no difference in gonadal-somatic index (GSI), an established biological metric of reproductive status and maturity, between F_1_ males behaving as ‘socially rock’ vs. ‘socially sand.’ (SI Figure 5). Our observation of divergent behavior between F_1_ brothers in the same tank suggests an interaction between the genome and the social environment.

**Figure 3:**
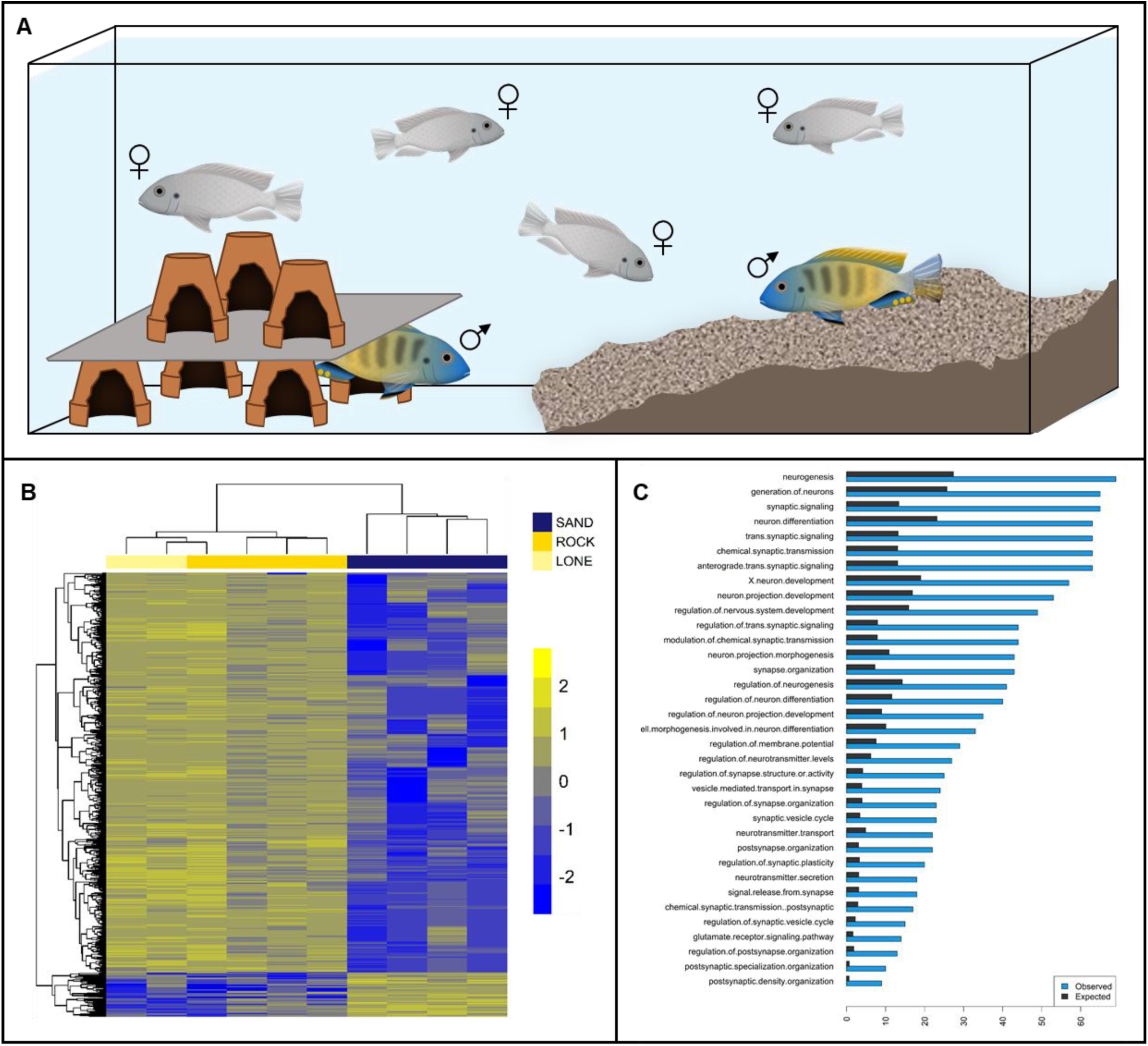
Genomics of divergent social context. (A) Schematic of the social context behavioral paradigm, in which rock-, sand- and rock-X sand-F_1_ hybrids were evaluated. (B) Heatmap of genes differentially expressed in the brains of F_1_ males behaving in either rock or sand social contexts. Each row in the heatmap is a gene, each column is an individual. Two F_1_ males were not paired with other males, and courted females over the rock habitat (lone). All other F_1_ males (n=8) were introduced to the testing arena in dyads. Notably, male brain gene expression clusters by social context and not fraternal relationships. (C) Gene Ontology (GO) Biological Process terms that show greater than expected overlap between (i) genes differentially expressed in the brains of social rock- vs. social sand-males and (ii) genes that are differentiated in the genomes of rock- vs sand-species groups.

We used RNA-seq to investigate gene expression profiles associated with behavior of rock-x sand-F_1_ hybrid males that were actively courting females over rock vs. sand. Whole brains of F_1_ males tested singly (n=2 lone) as well as F_1_ brothers assayed in dyads (n=4 dyads) were collected during courtship, and interrogated by RNA-seq. Strikingly, gene expression profiles clustered not by fraternal relatedness, but rather by behavioral context (Figure 3B). Males from dyads that courted females over rocks had expression profiles similar to single males (who also courted over rocks) but distinct from their brothers that built bowers and courted females over sand in the same tank. Genes were considered significantly differentially expressed between ‘social rock’ and ‘social sand’ brains if they exhibited both a fold change ≥ 2 and crossed the threshold of p_adj_ < 0.05. Based on this criterion, we found 832 genes differentially expressed between rock- vs. sand-behaving males (Figure 3B, SI Table 4). Among differentially expressed genes, we observed significant functional enrichment for GO Biological Process categories ‘synaptic signaling’ (p < 2.3e-21), ‘synaptic plasticity’ (p < 3.6e-09), ‘visual behavior’ (p < 2.09e-06); mouse phenotypes ‘abnormal learning/memory/conditioning’ (p < 5.9e-07), ‘abnormal telencephalon morphology’ (p < 3.95e-07), ‘abnormal spatial learning’ (p < 9.9e-07) and pathways ‘axon guidance’ (p < 3.3e-05), ‘oxytocin signaling’ (p < 5.2e-05) and ‘estrogen signaling’ (p < 1.8e-04) (SI Table 4). Matches against the Allen Brain Atlas database of gene expression yielded enrichment for exclusively sub-regions of the telencephalon: CA3, hippocampus (p_adj_ = 4.6e-6), CA2, hippocampus (p_adj_ = 7.3e-5), CA4, hippocampus (p_adj_ = 0.001), claustrum (p_adj_ = 0.001), subiculum (p_adj_ = 0.003), dentate gyrus (p_adj_ = 0.03) and the basomedial nucleus of the amygdala (p_adj_ = 0.03). The hippocampus encodes episodic memory and spatial representations of the environment (Olton, Becker et al. 1979), and more recently its subregions have been shown to play critical roles in anxiety, social interaction, and social memory formation (Hitti and Siegelbaum 2014, Zou, Chen et al. 2016, Chiang, Huang et al. 2018). Roughly 38% of differentially expressed genes also contained genetically differentiated SNP/InDels between rock- and sand-species (p-value < 2e-6, Fisher’s exact test), implying considerable cis-acting genetic variation. Enrichment of categories related to brain function and synaptic plasticity showed greater overlap than expected (Figure 3C). These context-dependent differences suggest rapid and concerted changes in brain gene expression as males experienced and responded to different social challenges and opportunities (O’Connell and Hofmann 2012, York, Patil et al. 2018).

## Discussion

### Genome-enabled inference of evolutionary change in morphology and behaviour

A fundamental problem in evolutionary biology is understanding the cellular, developmental and genetic basis of how traits change. This is a challenge because we lack sufficient information about how genes work in outbred genomes from nature and we do not fully comprehend the causal role of noncoding variation in specifying form and function. This problem is especially difficult for traits that are only observed in particular contexts, like development and behaviour. To make progress, we and others have focused on study systems exhibiting abundant phenotypic diversity built from a relatively small number of genetic changes. Here we identify and characterize the genetic variants that demarcate one of the deepest evolutionary splits amongst Lake Malawi cichlid groups, that between rock- and sand-dwelling species thought to have diverged in the last one million years. We found a small percentage (less than 0.1%) of genetic variants to be differentiated between rock- and sand-groups, and that the majority of differentiated variants (>96%) were noncoding. Differentiated non-coding variants were more likely to be in an evolutionarily conserved locus as a function of genetic differentiation, suggesting that divergent rock- vs. sand-noncoding changes are functional. To support this idea, we identified alternately fixed rock- vs. sand-noncoding variants within experimentally verified, vertebrate-conserved craniofacial and cranial neural crest cell enhancers. The latter observation is similar in type to the discovery of human-specific deletions within mammal-conserved regulatory sequence (McLean, Reno et al. 2011).

Recently we surveyed genome-wide divergence between sand-dweller sub-groups that construct pit vs. castle bowers, sand-made structures to attract females for mating (York, Patil et al. 2018). Mapping those variants to the same genome reference, we expected distinct patterns of diversification because rock-sand and pit-castle divergence likely occurred at different times, along different trait axes, under the control of different evolutionary forces (Streelman and Danley 2003). Consistent with expectation, there is clear clustering of genome divergence on chromosome 11 for the pit-castle comparison (SI Figure 6), while all chromosomes carry the signature of rock-sand diversification (Figure 1B). However, contrary to our expectation, rock- vs. sand- and pit- vs. castle-radiations have diverged in similar gene sets. Out of 3070 genes identified near 10kb high F_ST_ regions in the rock- vs. sand-comparison, 483 overlap with 1090 genes identified near high F_ST_ regions in the pit- vs. castle-comparison (p-value < 2e-9, Fisher’s exact test, SI Table 3). This result may imply that evolutionary diversification in Lake Malawi is limited, or constrained, by chromosomal location.

Overall, genes in proximity to rock-sand divergent variants were enriched for functional categories related to early forebrain and craniofacial development, neuronal function and social behavior. This list of variants, coupled with consistent patterns of functional and pathway enrichment, motivated follow up experiments focused on early brain development and adult social behavior (Figure 4). It is apparent from our work here and previously (Sylvester, Rich et al. 2010, Sylvester, Rich et al. 2013), that Malawi cichlid brains and nervous systems begin to differ during gastrulation in pathways that can be predicted from divergent genome sequences. This is interesting for at least two reasons. First, this observation runs counter to the ‘late equals large’ textbook example (Finlay and Darlington 1995) of how brains evolve differences in relative proportions of their parts (Sylvester, Rich et al. 2010). Similarly, such early variation in development is not thought to be a driving force in evolution, precisely because early changes can have global and ramifying effects. Collectively, our findings provide a partial description of the conditions wherein variation during the earliest stages of development can contribute to evolutionary diversification. In each case we have examined, variation in gene expression is quantitative, heterochronic and limited to a precise stage or time period.

**Figure 4:**
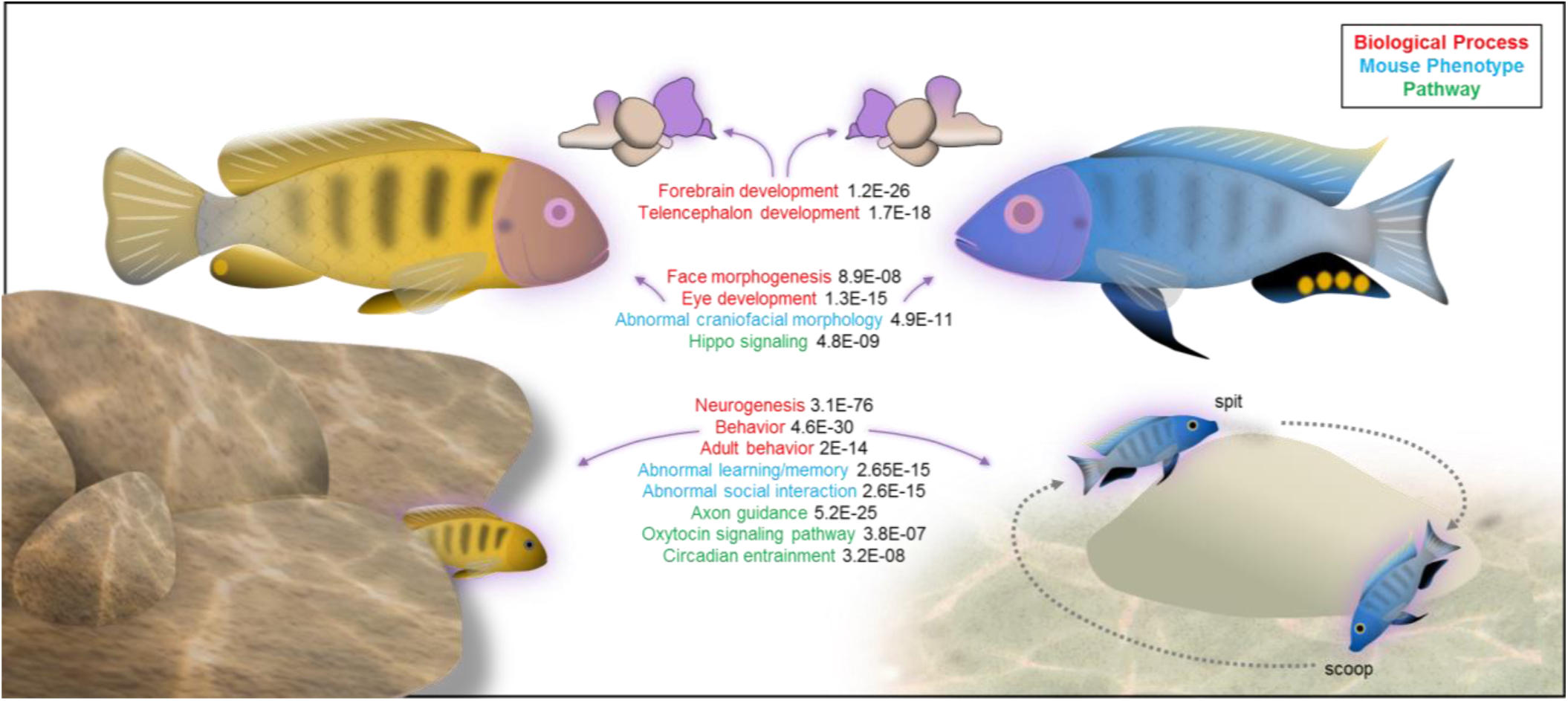
Genome-enabled discovery of evolutionary change in morphology and behavior. Summary cartoon synthesizing significant enrichment categories that differentiate the genomes of rock- vs. sand-dwelling Malawi cichlids. Strong and consistent enrichment of craniofacial, neural and behavioral categories motivated follow on experiments in early brain development (Figure 2) and adult social behavior (Figure 3).

Sydney Brenner recognized the relationship between the genetic specification of nervous systems and the behavioral output of the brain (Brenner 1974). However, because these events take place so far apart in the lifespan of a vertebrate, they are rarely studied simultaneously. Here, the genome connects the two phenomena: rock- vs. sand-divergent gene sets indicating that both brain development and social behavior have been under divergent selection during the evolutionary diversification of these groups. To evaluate social behavior in rock- vs. sand-dwelling Malawi cichlids, we constructed a social context arena. The presence of sand and simulated rocky caves was sufficient to elicit species-appropriate male behavior when rock- or sand-males were tested with rock- or sand-gravid females. When rock-x sand-F_1_ hybrid males were tested, one per tank, with F_1_ hybrid females, males courted females in the rock quadrant of the tank. Notably, when dyads of F_1_ brothers were tested in this tank paradigm with gravid F_1_ females, we observed simultaneous ‘social rock’ and ‘social sand’ behavior. Brain gene expression profiles from behaving males clustered by social context (social rock vs social sand), and not by fraternal relationships. Differentially expressed genes were enriched for brain regions and pathways implicated in social interaction and overlapped significantly with rock- vs. sand-divergent genetic variants.

Social context is known to influence the brain. For instance, our clustering results are similar to those of Whitfield and colleagues (Whitfield et al. 2003) who showed that brain gene expression in honey bees was predictive of behavior. Likewise, changes in brain morphology and gene expression predictably accompany the ascent to dominance in the cichlid fish *Astatotilapia burtoni* (Fernald and Maruska 2012). Our data seem not to fit the model of dominant-subordinate however. In our experiments, the gonado-somatic index (GSI) did not differ between social rock vs. social sand brothers within dyads. Both males exhibited nuptial coloration, courted females and in cases with multiple gravid females, both brothers reproduced. Body size was associated with divergent social rock vs. social sand behavior of F_1_ males; the social rock brother was always larger (mean mass was 26.96g ± 3.4 [SE] compared to 19.45g ± 2.2).

Our experiments demonstrate that F_1_ hybrid male brains can express both social rock- and social sand-behavioral programs, and that social context determines which program is executed. This observation is similar to, but also different than, pit-digging x castle-building F_1_ male Malawi cichlids who carry out parental bower behaviors in a specific sequence (York, Patil et al. 2018). Notably, in both cases, the hybrid males exhibit one of the two different parental behaviors at any one time – there is no intermediate behavior. In the pit- vs. castle-case, we think that the bower structure itself and/or a threshold signal from females might lock the hybrid male brain into a behavioral state. In the rock- vs sand-case here, it appears that other social cues (i.e., the presence and size of a rival male) lock the hybrid male brain into a behavioral state. These context-dependent behaviors, accompanied by changes in brain gene expression, are compelling examples of interaction between the genome and the social environment. The cellular and genetic basis of these behaviors and their plasticity deserves further attention. Our comparative genomic and brain gene expression data, combined with enrichment testing and experimental approaches, highlight that the Malawi cichlid telencephalon will be central to this future work.

## Supporting information

Supplementary Table 1 Species and seq characteristics

Supplementary Table 2 genomewide identity of enrichment - topfun - SFARI - enhancers

Supplementary Table 3 RNA-seq DEG

Supplementary Table 4 rocksand versus pitcastle

Supplementary Table 5 Samples for RNA seq

## Acknowledgements

This work was supported by grants from the NIH (R01GM101095, 2R01DE019637-10 to JTS and F32GM128346-01A1 to ZVJ) and the Human Frontiers Science Program (RGP0052/2019 to JTS). We would like to thank Shweta Biliya and the Genomics Core at Georgia Tech for invaluable assistance with the NGS sequencing. We also thank the members of the Streelman lab for comments on this manuscript.

## Competing interests

The authors declare no competing interests.

## Methods

### Genome sequencing

We extracted genomic DNA, from fin clips (Qiagen DNeasy, Cat #69504), from 8 rock dwelling and 14 sand dwelling Lake Malawi species (SI Table 2). We made libraries using the Illumina Nextera Library prep kit and performed paired-end sequencing on the Illumina Hi-Seq 2500 at Georgia Tech. The *Metriaclima zebra* reference genome version MZ_UMD2a (Conte, Joshi et al. 2019) was used for genome alignment, variant discovery and annotation using standard BWA (Li and Durbin 2009) and GATK practices (Van der Auwera, Carneiro et al. 2013). The maximum likelihood tree in Figure 1A was constructed using SNPhylo (Lee, Guo et al. 2014), from variant data.

### Genetically Divergent Regions

Vcftools (Danecek, Auton et al. 2011) was used to calculate F_ST_ (--weir-fst-pop) between the 8 rock and 14 sand species. Variants with F_ST_ = 1 were noted to be alternately fixed between rock and sand lineages in our dataset. F_ST_ was also measured across 10kb windows (--fst-window-size). Significance thresholds were marked using the fdrtool package in R. All variants were annotated using Snpeff 4.3i (Cingolani, Platts et al. 2012). We tested the genes within 25 kb of significantly differentiated variants for enrichment of functional categories. The cichlid gene names were converted to human analogs using Treefam based mapping (Ramakrishnan Varadarajan, Mopuri et al. 2018) and functional enrichment was determined using the TOPPFUN web-browser interface (Chen, Bardes et al. 2009).

### PhastCons analysis

Pairwise alignments were generated using lastz v1.02(Harris 2007), with the following parameters: “B=2 C=0 E=150 H=0 K=4500 L=3000 M=254 O=600 Q=human_chimp.v2.q T=2 Y=15000”. This was followed by using USCS genome utilities (https://genome.ucsc.edu/util.html, https://hgdownload.soe.ucsc.edu/admin/exe/linux.x86_64/FOOTER) axtChain tool with - minScore=5000. Additional tools with default parameters were then used following the UCSC whole-genome alignment paradigm (http://genomewiki.ucsc.edu/index.php/Whole_genome_alignment_howto) in order to obtain a contiguous pairwise alignment. Multiple alignments were generated from pairwise alignments with the multiz v11.2 (Blanchette, Kent et al. 2004) program, using default parameters and the following pre-determined phylogenetic tree: ((((*M. zebra, P. nyererei*), *A. burtoni*), *N. brichardi*), *O. niloticus*) in agreement with Brawand et al. (Brawand, Wagner et al. 2014). Sequence conservation scores were then obtained using PhastCons (Siepel, Bejerano et al. 2005) with a phylogenetic model estimated by the phyloFit (Siepel and Haussler 2004) program, both from the PHAST software package (v.1.3). The model fitting was done using default parameters. PhastCons was run in two iterations, first to obtain the free parameters of the model (--estimate-trees and --no-post-probs) and then using the output from this we ran PhastCons again to attain the conservation scores with --target-coverage 0.3 --expected-length 100.

### Vertebrate-conserved enhancer elements

A comparative genomic approach was used to identify putative craniofacial and neural crest CNEs in mammals that segregate SNPs between rock-sand cichlid species. Experimentally verified and published genome-wide craniofacial and neural crest enhancers active during early embryonic stages that play a role in shaping the development of neural crest and craniofacial structures in mammals were identified from published literature (Rada-Iglesias, Bajpai et al. 2012, Attanasio, Nord et al. 2013). We used the liftOver tool (Kent, Sugnet et al. 2002), which maps orthologous genomic regions between species to convert genomic coordinates from one species to another. Using a Human to *Oreochromis niloticus* to *Metriaclima zebra* mapping and a Mouse to *Oreochromis niloticus* to *Metriaclima zebra* mapping, we identified the orthologous genomic locations of the published craniofacial and neural crest enhancers in cichlids. We designated any alternately fixed variant (variant with F_ST_ = 1) that was also within an orthologous CNE as putatively involved in the rock-sand divergence (SI Table 2).

### Brain region enrichment analysis

We identified 10,391 cichlid genes with human homologues and generated an expression matrix for each gene across 250 human brain structures spanning telencephalon, diencephalon, mesencephalon, and metencephalon using adult human brain microarray data collected by the Allen Brain Institute (Hawrylycz, Lein et al. 2012). Cortical regions and gyri for which fish do not have putative homologues were excluded from the analysis (100/350, leaving 250 regions for subsequent analysis). The expression matrix was generated using the get_expression function in the ABAEnrichment Bioconductor package in R (Grote, Prufer et al. 2016). We then calculated the specificity of expression for each gene in each of these brain regions using the specificity.index function in the pSI package for R. This function calculates a matrix of gene expression specificity indices, and corresponding p-values, as described previously (Dougherty, Schmidt et al. 2010, Xu, Wells et al. 2014). We then tested whether 1) genes within 25kb of rock vs. sand significantly differentiated variants (described above under “Genetically Divergent Regions”), and 2) genes that were differentially expressed between rock- vs. sand-behaving F1 hybrid males, were enriched for transcriptional markers of specific brain regions using the *fisher*.*iteration* function with Benjamini-Hochberg correction, again using the pSI package for R. For enrichment testing of differentially expressed genes, we restricted analysis to genes that met the following criteria: 1) transcripts for the gene were detected in all eight paired behaving males, and 2) at least 6 transcripts were detected in each subject.

### Staging during gastrulation

Cichlid gastrulation was split into three sub stages within the gastrula stage 9 (Murata, Tamura et al. 2010). Gastrulation lasts 8 to 12 hours, depending on the species, and is defined as after the shield (as described in zebrafish) stage until the presence of the first somite at the beginning of neurula (stage 10). Embryos were classified as early gastrula (EG) by an asymmetry in epiboly after shield stage until the formation of a ridge that is analogous to the anterior neural ridge (ANR) in chick and mouse and the anterior neural border (ANB) in zebrafish. At that point embryos were classified as mid gastrula (MG). MG lasts until the formation of the dorsal-ventral axis, defined by further lengthening of one side of the embryo, which begins to thicken as epiboly progresses. This is the dorsal side of the embryo, and the side opposite the ANR is classified the ventral side of the embryo. At this point the embryos are defined as late gastrula (LG). LG ends with the specification of the neural plate, which appears as a portion of the dorsal embryo that is raised relative to ventral side, usually in line with the ANR.

### Immunohistochemical staining

Embryos were harvested at 24 hours post fertilization (hpf) from each of the rock -dwelling cichlids *Metriaclima patricki* and *Metriaclima zebra* and the sand-dwelling cichlid *Copadichromis borleyi* and *Tramitichromis intermedius*. The embryos were cultured until they reached gastrula stage, approximately 36 to 40 hpf, then fixed at intervals throughout gastrula until neurula. The embryos were then treated with auto-fluorescence reducer (1.55mL 5M NaCl, 250ul Tris-HCl, pH 7.5, and 95mg NaBH4) overnight, and 10% 2-mercaptoethanol for 1 hour. Next, whole mount *in situ* hybridization was done, using a modification methods we published previously (Fraser, Bloomquist et al. 2008). *irx1b* was visualized using Fast Red (naphthol chromogen, Roche Diagnostics), which fluoresces at near red wavelengths (500-650 nm). After *in situ* hybridization, embryos were immunostained for pSMAD 1,5,8 protein, using published protocols (Tucker, Mintzer et al. 2008). Embryos were then bathed in Vectashield (Vector Labs) containing DAPI and placed in a specially built mold(White, Sylvester et al. 2015) that accommodates the large yolk and holds the embryo upright. Embryos were then scanned using a Zeiss LSM 700-405 confocal microscope and processed using LSM 700 software and Image J.

### Rock-Sand hybridization and genotyping

Two rock-sand crosses, one between *Copadichromis borleyi* (CB, sand-dweller sire) and *Metriaclima zebra* (MZ, rock-dweller dam) and another between *Mchenga conophoros* (MC, sand-sire) and *Petrotilapia sp*. ‘thick bar’ (PT, rock-dam), were artificially generated by taking the eggs from the dam just prior to spawning and mixing with sperm from the sire. The resultant F_1_ were grown in tanks and allowed to spawn normally to generate F_2_. Several F_2_ broods were taken from multiple F_1_ females for each cross, a total of 355 individuals for the CB x MZ cross and 608 for the MC x PT cross. The embryos were fixed at every stage starting at gastrula (stage 9) until early pharyngula (stage 14). The F_2_ embryos were RNA-extracted at stage 9. DNA extraction was performed by fixing the embryos (stage 11-14) in 70% ethanol, then removing the tail from each individual and extracting the DNA using an extraction kit (Qiagen). Following extraction, the F_2_ embryos were genotyped using custom probes (CAAATCTCCC[C/T]CCGCGGC, Taqman custom probes, Invitrogen) designed to identify a SNP in *irx1b* using RT-PCR. A subset of the embryos was also sequenced at a 900 bp interval around the *irx1b* SNP to verify the custom probes.

### Quantitative F_2_ Analysis

We quantified *irx1b* in F_2_ at stage 9 and separated by genotypic class. The 74 heterozygous rock X sand F_2_ embryos were dissected to remove most of the yolk and the total RNA was extracted from each individual using an RNA Extraction Kit (Qiagen). The amount of mRNA specific to each allele of *irx1b* was quantified by using the RNA-to-Ct kit (Invitrogen) and the custom probes. The delta Ct for each heterozygote was generated with the equation, 2^(allele from dam – allele from sire). We tested the data with an ANOVA, followed by a Tukey’s multiple comparison test to determine significance between genotype classes.

### Forebrain and eye measurements

The forebrain and eyes were measured by integrating the area of transverse sections in embryos of rock- and sand-dweller cichlid species, using previously published methods (Sylvester, Rich et al. 2013). The rock-dweller species included *Cynotilapia afra* (CA, planktivore), *Labeotropheus fuelleborni* (LF, algivore) and *Metriaclima zebra* (MZ, generalist); sand-dweller species included *Aulonocara jacobfreibergi* (AJ, ‘sonar’ hunter), *Copadichromis borleyi* (CB, planktivore) and *Mchenga conophoros* (MC, insectivore/generalist). Embryos from each species, as well as the F_2_ individuals, were measured starting from the earliest the telencephalon can be differentiated from the forebrain (mid-somitogenesis, stage 12) and at each subsequent stage until the forebrain has defined prosomeres (early pharyngula, stage 14) (Sylvester, Rich et al. 2010). To keep measurements standardized across stages, all measurements were defined by forebrain morphology at the earliest timepoint (stage 12). The ‘eye’ measurement remains consistent at all stages, the ‘anterior’ measurement includes the telencephalon and presumptive olfactory bulb, and the ‘posterior’ measurement includes the diencephalon and each of its constitutive prosomeres (dorsal and ventral thalamus and hypothalamus). To facilitate measurements, we used gene expression of *rx3* (for stage 12 embryos) and *pax6* (stage 13 and 14) to identify the different structures of the forebrain and eye.

### RNA Extraction and Sequencing, Adult Social Behavior

Adult F_1_ hybrid males (Supplementary table 4) were introduced to an assay tank containing females of the same cross and simulated rock habitat on one side and simulated sand habitat on the other side separated by empty tank space (Figure 3A). Male brains were harvested within 20 minutes of exhibiting territoriality and displays for females by rapid decapitation and whole brains were immediately stored in RNAlater (Thermo Fisher Cat# AM7020).

Tissues were frozen in liquid nitrogen, homogenized using a mortar and pestle and placed in trizol. Following standard chloroform extraction, RNeasy mini columns (Qiagen Cat No./ID: 74104) were utilized to purify RNA for sequencing. Total RNA was quantified using Qubit (Molecular Probes) and quality analyzed using the Agilent 2100 Bioanalyzer System for RNA library preparation. RNA input was normalized to 1µg and libraries were prepared using the TruSeq Stranded mRNA Sample Prep Kit (Illumina-Kit A). Libraries were again quantified, quality assessed, and normalized for sequencing on the HiSeq 2500 Illumina Sequencing System (Georgia Tech Genomics Core, standard practices). Experimental design and raw files can be accessed on the NCBI Gene Expression Omnibus database under the accession number GSE122500.

### Differential Gene Expression Analysis

Raw sequence reads from whole brain transcriptomes were quality controlled using the NGS QC Toolkit (Patel and Jain 2012). Raw reads with an average PHRED quality score below 20 were filtered out. Filtered reads were also trimmed of low-quality bases at the 3’ end. High quality sequence reads were aligned to the *M*.*zebra* reference genome MZ_UMD2a(Conte, Joshi et al. 2019) using TopHat v2.0.9 (Trapnell, Pachter et al. 2009). On average, across all samples, over 95% of reads mapped to the reference genome. The resulting TopHat2 output bam files were sorted and converted to sam files using samtools v0.19 (Li, Handsaker et al. 2009). Sorted sam files were used as input for the HTSeq-count v0.6.1 program to obtain fragment counts for each locus (Anders, Pyl et al. 2015). Fragment counts were scale-normalized across all samples using the calcNormFactors function in the edgeR package v3.6.8 (Robinson, McCarthy et al. 2010). Relative consistency among replicates and samples was determined via the Multidimensional scaling (MDS) feature within the edgeR package in R. The native R function *hclust(dist)* used to cluster samples. Scale-normalized fragment counts were converted into log_2_ counts per million reads mapped (cpm) with precision weights using voom and fit to a linear model using limma v3.20.9 (Ritchie, Phipson et al. 2015). Pairwise contrasts were constructed between socially rock and socially sand samples. After correcting for multiple comparisons using the Benjamini-Hochberg method (Hochberg and Benjamini 1990), genes were considered differentially expressed between socially rock and socially sand samples if they exhibited both a fold change ≥ 2 and P_adj_ < 0.05. Using Treefam based mapping (Ramakrishnan Varadarajan, Mopuri et al. 2018) the cichlid gene names were converted to human analogs and functional enrichment was determined using the TOPPFUN web-browser (Chen, Bardes et al. 2009).

---

**Table 5.**
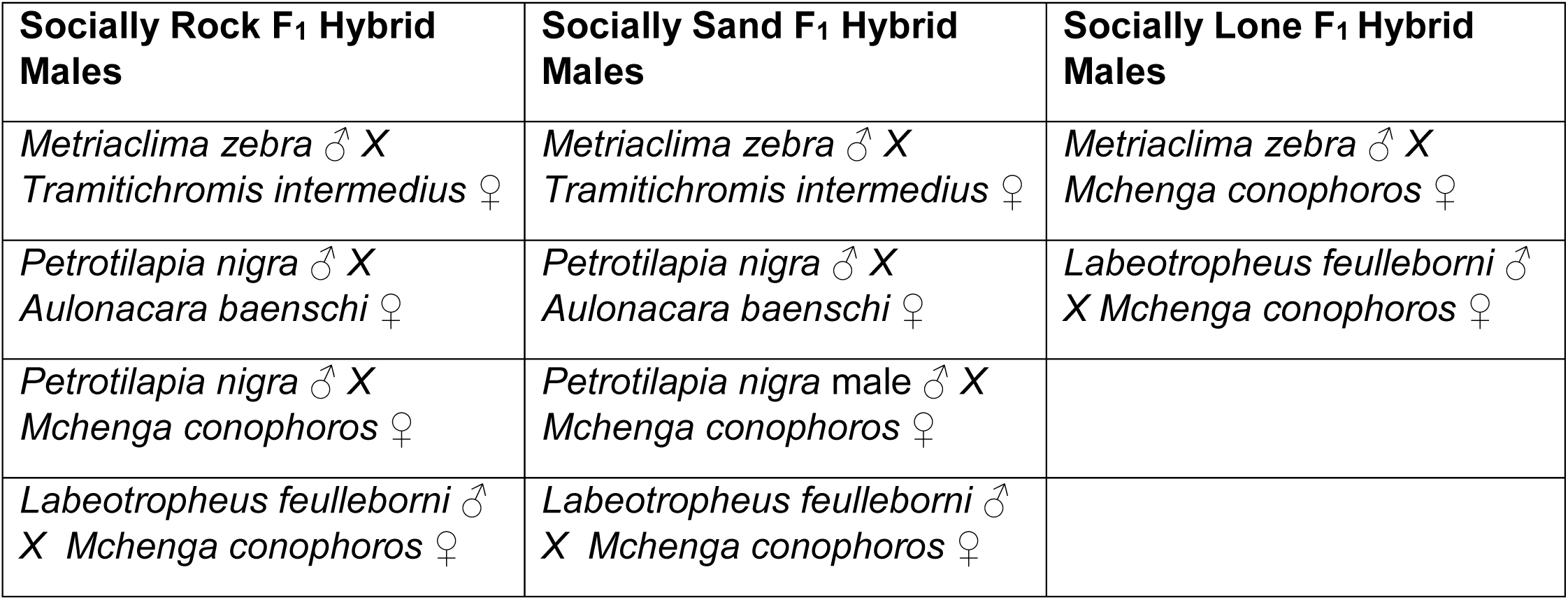

**SI Figure 1:**
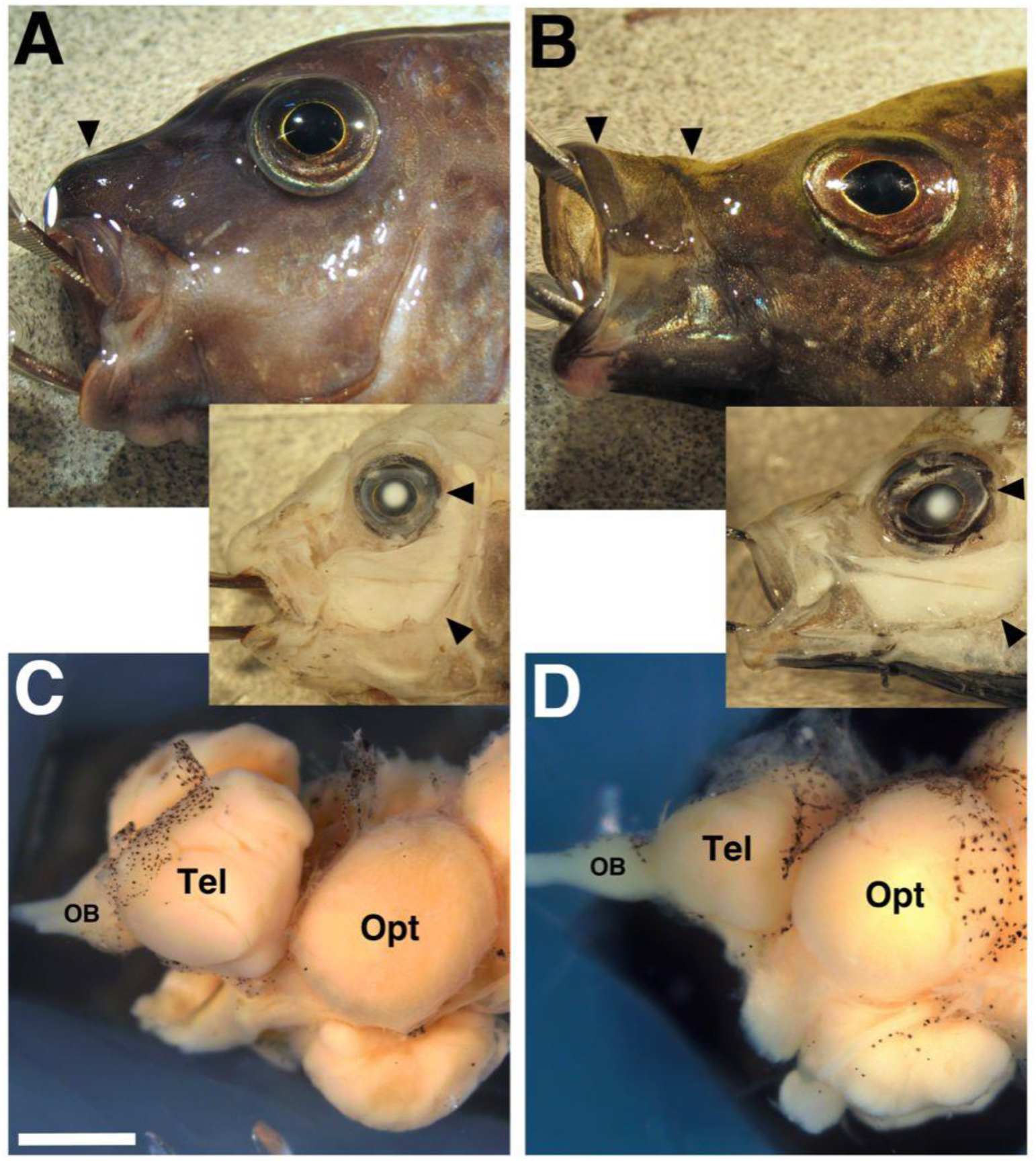
Differences in the face (A-B) and brain (C-D) between rock- vs. sand-dwelling Lake Malawi cichlids. Rock-dwellers (A, C) have strongly reinforced jaws, smaller eyes, typically larger cheek muscles, larger olfactory bulbs and telencephala. Sand-dwellers have kinematic, gracile jaws, larger eyes, less robust cheek musculature, and large optic tecta.

**SI Figure 2:**
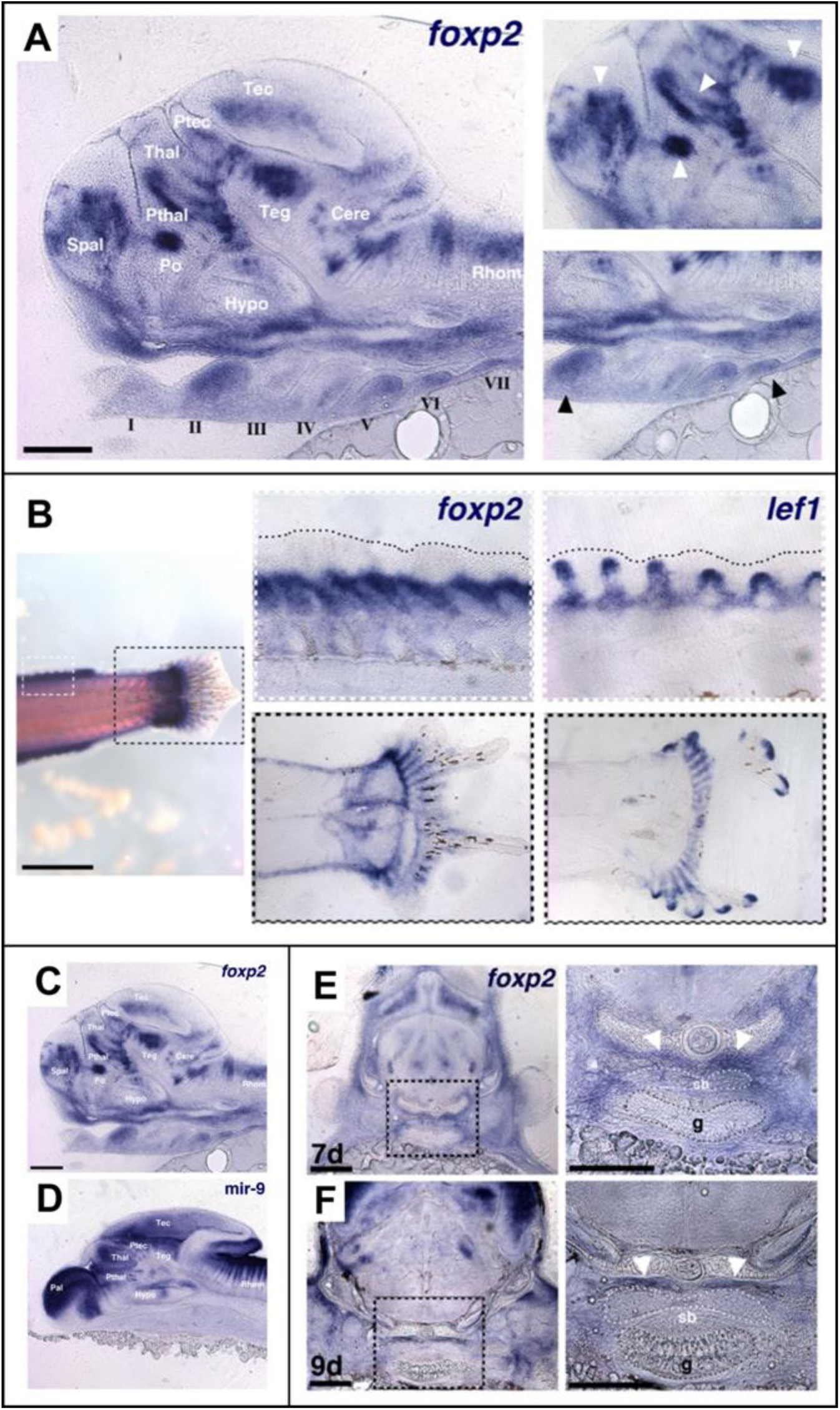
Malawi cichlid *foxp2* is expressed in the brain and in all sonic organs. We previously noted novel expression domains for cichlid *foxp2* (Bloomquist, Fowler et al. 2017), elaborated here (A) Expression of *foxp2* throughout the brain (regions labeled in white) and developing pharyngeal arches (labeled I – VII) at 5 days post fertilization (dpf). On the far right, white arrows indicate expression in developing ganglia in the midbrain, diencephalon, pre-optic region, and subpallium. Black arrows point to expression in pharyngeal arches II and V. (B) *foxp2* expression, along with the WNT pathway transcription factor *lef1*, in the developing dorsal and tail fins at 7 dpf. The white- and black-dashed boxes on the right-most panels are zoomed in, midline sections of the boxes on the left. Adjacent, overlapping expression of *foxp2* and *lef1* are indicative of interaction between *foxp2* and the WNT pathway (Bonkowsky, Wang et al. 2008). (C) and (D) show the expression of *foxp2* and *micro RNA-9* (*mir-9*). *mir-9* has been shown to regulate *foxp2* activity in vertebrates (Shi, Luo et al. 2013) and the anti-correlated expression patterns in cichlids suggest a similar interaction. (E) and (F) document expression of *foxp2* in the developing swim bladder at 7 and 9 dpf. The black-dashed boxed on the left panels indicate the zoomed panels on the right. *foxp2* is generally expressed in the mesenchyme within and dorsal to the swim bladder at 7 dpf (white arrows). Once the swim bladder epithelium forms by 9 dpf, *foxp2* expression is localized dorsally (white arrows). All scale bars are 100μm. Abbreviations: Rhom = Rhombencephalon, Cere = Cerebellum, Teg = Tegmentum, Tec = Optic Tectum, Ptec = Pretectum, Thal = Thalamus, Pthal = Prethalamus, Po = Pre-optic area, Hypo = Hypothalamus, Spal = Subpallium, sb = swim bladder, g = gut.

**SI Figure 3:**
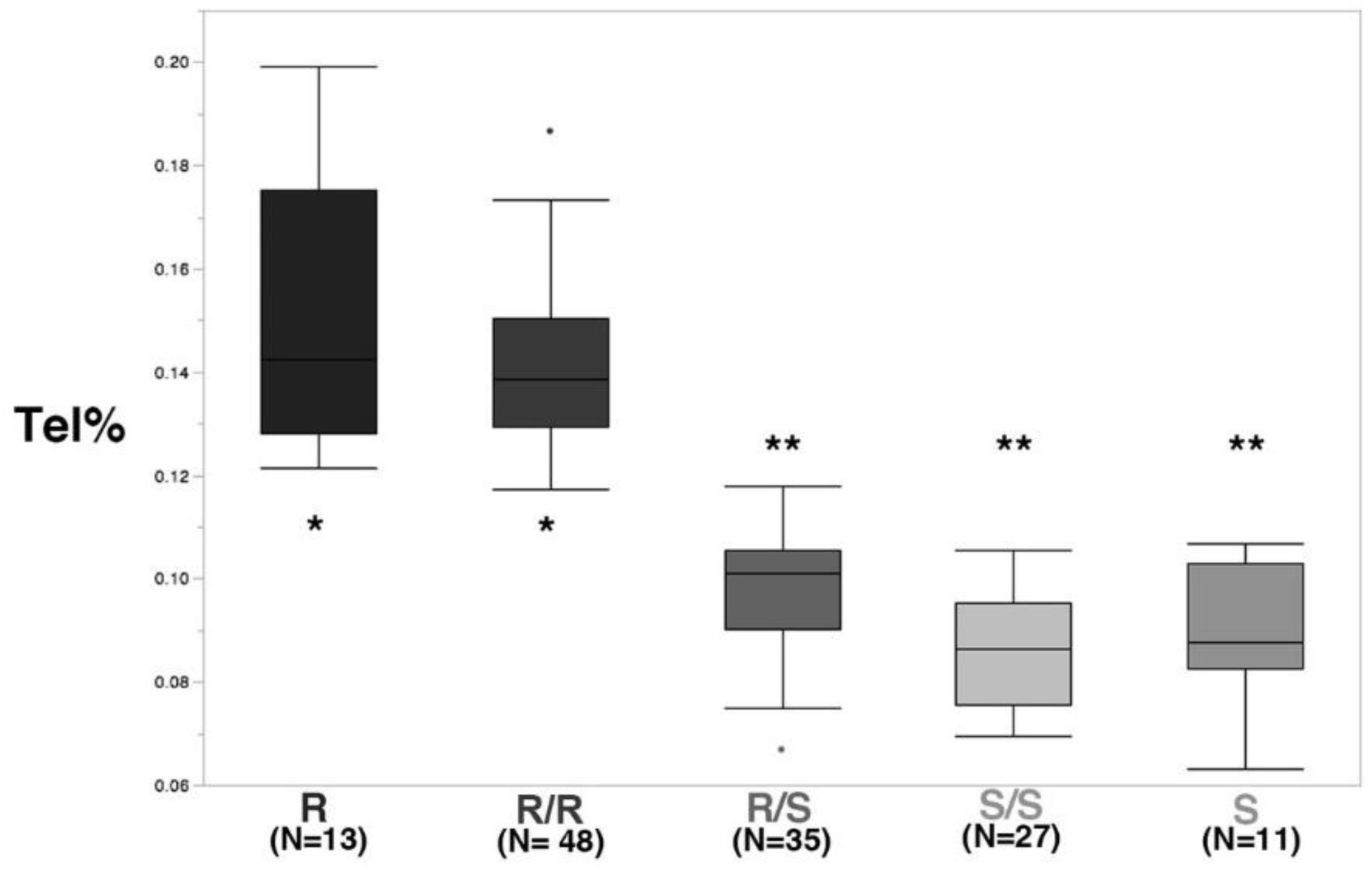
The relative size of the telencephalon differs in rock-, sand- and F_2_ hybrids indexed for *irx1b* genotypes. Rock-, sand- and F_2_ hybrid individuals indexed for *irx1b* genotype were sampled at stages 12-14. We calculated the volume of the telencephalon in each individual and express this as a percentage of total forebrain volume. See also Figure 2.

**SI Figure 4:**
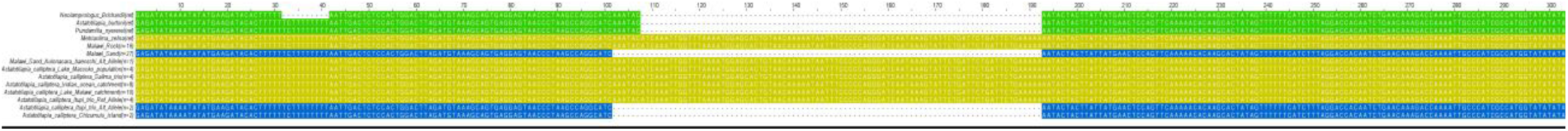
Schematic of an InDel located in the 3’UTR of the Malawi cichlid *irx1b* gene. Summary of allelic states: (1) rock-dwelling species possess an 85bp insertion (yellow) with similarity to Rex1 non-LTR retrotransposon, compared to outgroup species (shown in green); (2) sand-dwellers generally lack the insertion and exhibit a 6bp deletion (blue), compared to outgroup species. Note that *Aulonocara baenschi* is heterozygous. Most individuals of *Astatotilapia calliptera* carry the rock-insertion allele; however, individuals at Chizumulu Island and Itupi possess the sand allele.

**SI Figure 5:**
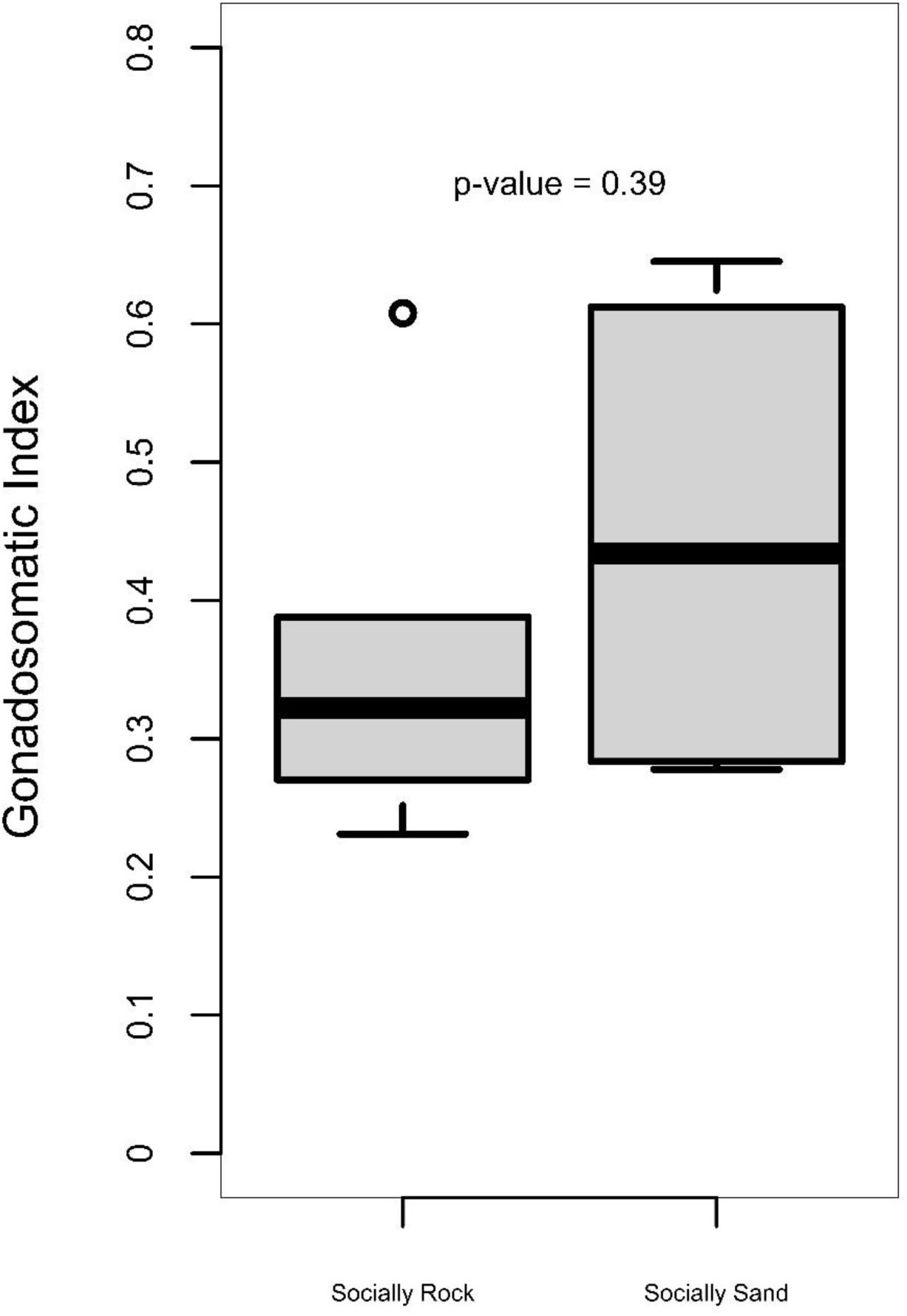
Gonadosomatic index (GSI) does not differ amongst males behaving as social rock- or social sand-. See also Figure 3.

**SI Figure 6:**
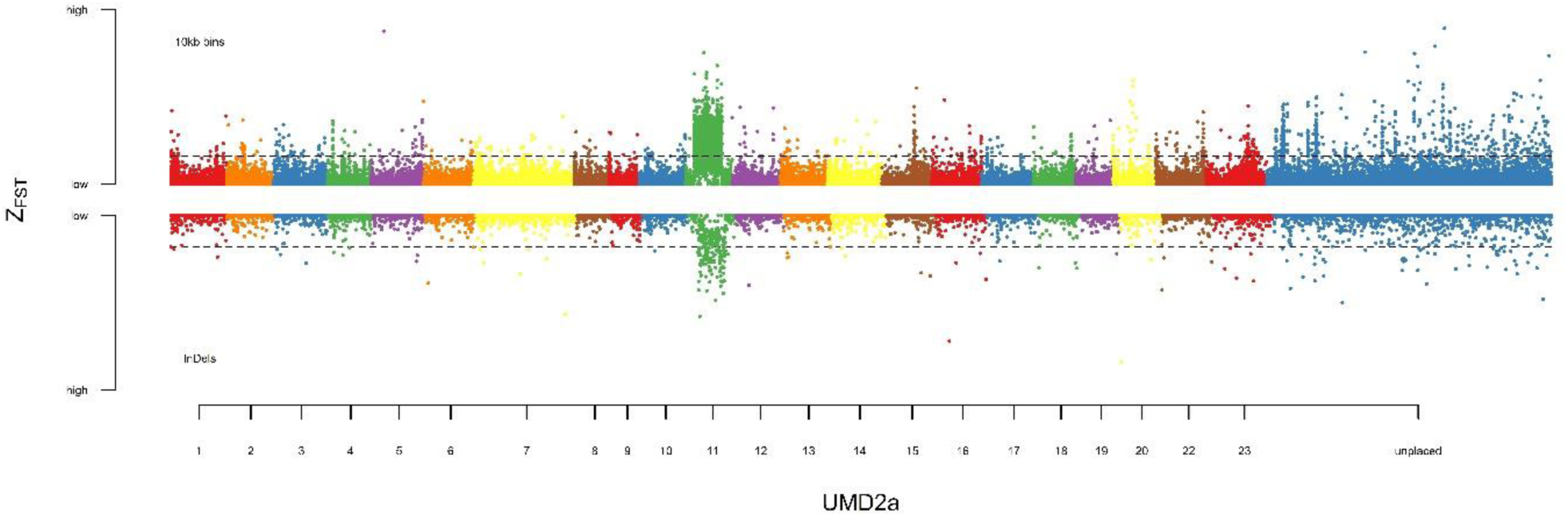
Genomic differentiation amongst sand-dwelling species that construct pit vs. castle sand bowers. Z-F_ST_ plot shows genome divergence amongst pit-digging vs. castle-building sand-dweller species, updated by mapping variants to the UMD2a reference genome (data from (York, Patil et al. 2018)). Threshold lines indicate 2.5% FDR.

## Notes

### Competing Interest Statement

The authors have declared no competing interest.

